# Nebula: Ultra-efficient mapping-free structural variant genotyper

**DOI:** 10.1101/566620

**Authors:** Parsoa Khorsand, Fereydoun Hormozdiari

## Abstract

**Motivation:** Large scale catalogs of common genetic variants (including indels and structural variants) are being created using data from second and third generation whole-genome sequencing technologies. However, the genotyping of these variants in newly sequenced samples is a nontrivial task that requires extensive computational resources. Furthermore, current approaches are mostly limited to only specific types of variants and are generally prone to various errors and ambiguities when genotyping events in repeat regions. Thus we are proposing an ultra-efficient approach for genotyping any type of structural variation that is not limited by the shortcomings and complexities of current mapping-based approaches.

**Results:** Our method Nebula utilizes the changes in the count of *k*-mers to predict the genotype of common structural variations. We have shown that not only Nebula is an order of magnitude faster than mapping based approaches for genotyping deletions and mobile-element insertions, but also has comparable accuracy to state-of-the-art approaches. Furthermore, Nebula is a generic framework not limited to any specific type of event.

**Availability:** Nebula is publicly available at https://github.com/Parsoa/NebulousSerendipity

## 1 Introduction

Structural variants (SVs) are defined as medium and large size genomic alterations. SVs have many different types, e.g. deletion, duplication, inversions and mobile-element insertions (Alkan et al. (2011)). It has become clear that SVs are a major contributing factor in human disease (Stankiewicz and Lupski (2010)) and evolution (Zhang et al. (2009)). However, efficient and accurate genotyping of the all types of SVs using whole-genome sequencing (WGS) data is not a trivial task. In many of the large scale genomic studies structural variants are being ignored or they are an afterthought. The main reason behind SVs not being as thoroughly studied as other types of variants such as SNVs, is due to complexity of accurate discovery and genotyping of these types of variants. This negligence in considering SVs is contributing to the missing heritability in complex disorders (Manolio et al. (2009); Eichler et al. (2010)).

With advent of high-throughput sequencing (HTS) technologies, we have made great progress in understanding the contribution of structural variants (SVs) in diseases and evolution. In the 1000 genomes (1KG) project a set of over 42,000 SVs were discovered and genotyped in 2,500 samples (Sudmant et al. (2015)). Recently, a few samples (e.g., CHM1 or CHM13) were sequenced using long-read technologies (i.e., PacBio). A comparison of the calls made using the state-of-the-art computational methods for SV discovery using the short-read HTS (e.g., LUMPY (Layer et al. (2014)), DELLY (Rausch et al. (2012)), pindel (Ye et al. (2009)), Tardis (Soylev et al. (2017)) against the SV calls produced using long-read options clearly indicates that many SVs (*>* 50%) are not being predicted by our best practices (Chaisson et al. (2015)). Thus, we are in need of approaches which can efficiently genotype these newly found SVs in the large number of WGS samples.

With additional samples being whole-genome sequenced at a breathtaking rate, an approach to accurately and efficiently genotype the (common) SVs in them is needed. In addition, with more comprehensive sets of SVs being predicted using long-read technologies we would like to be able to genotype them in the samples that have already been processed.

The current approaches for genotyping SVs using WGS data are mainly based on first mapping the reads to the reference genome and then predicting the genotype (Handsaker et al. (2011)). This framework has three main drawbacks. First, the mapping step is quite resource intensive. Second, these approaches are only limited to specific types of variants (SNV, small indels and large CNVs). Third, genotyping any variant close to repeats in the reference genome would be less accurate due to potential of inaccurate mapping.

### Mapping free approaches

Mapping-free approaches are becoming popular for different genomic and transcriptome applications. The mapping-free approaches are not limited by the shortcoming of the reference genome and tend to be more efficient. These approaches also have been recently utilized successfully in variant discovery with different biological applications. One of the first tools to introduce a mapping-free method for variant discovery is Cortex (Iqbal et al. (2012)). Cortex introduces the concept of colored de bruijn graph to compare the *k*-mers from different samples to predict variants between the samples (Iqbal et al. (2012)). Cortex was also used successfully for predicting variants in the 1000 genomes project. The methods DISCOSNP (Uricaru et al. (2014)) is able of predicting SNPs efficiently in multiple sequenced samples without using the reference genome. NovoBreak is one the tools which utilizes *k*-mer counts to predict somatic variants between tumor and normal whole-genome sequence samples, include structural variants (Chong et al. (2017)). Another application of mapping-free approaches in discovery of *de novo* variants in families. The tool Scalpel (Narzisi et al. (2014)) is a mapping-free approach for accurate discovery of *de novo* indels in whole-exome sequenced samples. Recently, mapping-free approaches have also been utilized in improving the association studies using whole-genome sequencing data (Rahman et al. (2018)). The tool HAWK (Rahman et al. (2018)) is capable of fast and accurate discovery of variants associate with phenotype of interest by comparing the *k*-mers frequencies between cases and controls. These growing application and tools for reference-free approaches has also resulted in development of several tools for fast and accurate *k*-mer quantification. Few of the tools used for fast *k*-mer quantification include JellyFish (Marçais and Kingsford (2011)), Khmer (Crusoe et al. (2015)), DSK (Rizk et al. (2013)) and KMC (Deorowicz et al. (2013)).

Here we are proposing a novel mapping-free approach, Nebula, which utilize *k*-mer counts for efficient and accurate prediction of genotypes (i.e., {0/0, 0/1, 1/1}) of (common) structural variant in any wholegenome sequenced (WGS) sample. The proposed method has two main stages of preprocssing and genotyping. In the preprocessing stage, a set of *k*-mers whose counts would drop or increase as a result of each SV are selected from samples affected by those SVs. During the genotyping stage, the selected *k*-mers are counted in new whole-genome sequencing sample and the variants are called based on differences between observed and expected *k*-mer counts.

## 2 Methods

Nebula models the problem of genotyping structural variation as that of measuring changes in *k*-mer counts in the newly sequenced samples. The method consists of two main phases: (i) *k*-mer preprocessing phase: Provided a set of SVs as input, Nebula selects a set of *k*-mers whose frequency is different in the alternate allele versus reference allele as a result of these SVs. These *k*-mers will be used in the next stage to genotype the same set of SVs on any large number of WGS samples. This step is preformed only once for any SV using the coordinates of the SV and the sequence data (WGS data or assembly) of the sample(s) that has the alternate alleles for these SVs. (ii) Genotyping phase: The main phase of the Nebula and the crux of the method is the genotyping stage. Provided the set of *k*-mers associated with the input SVs, Nebula will count these *k*-mers in the WGS samples for genotyping. These counts will then be plugged into a linear program to produce real-valued estimates for the copy number of these SVs in the newly sequenced sample. A final rounding converts the real genotype values to three states of 0/0, 0/1 or 1/1. A graphical representation of this pipeline can be seen in Figure 1.

**Figure 1:**
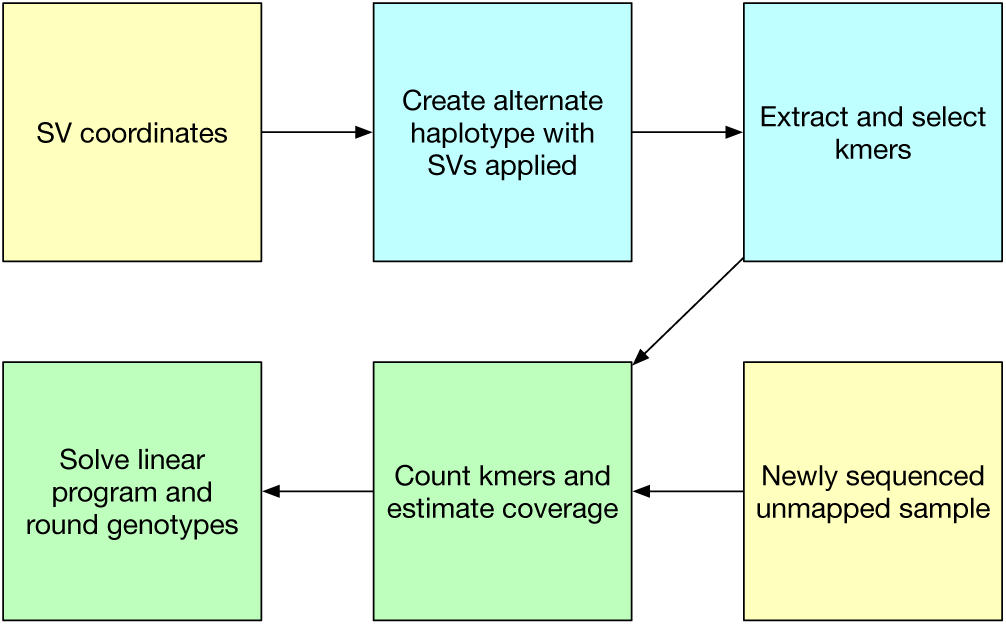
Simplified overview of Nebula’s pipeline. Yellow boxes show inputs while blue boxes represent the preprocessing stage and green boxes represent the genotyping stage.

Note that, the preprocessing phase is ran only once. After the appropriate *k*-mers are selected in the preprocessing phase, only the genotyping phase needs to be repeated for genotyping of the input/common SVs on any new WGS samples.

### 2.1 *k*-mer selection phase

The choice of *k*-mer length was made based on various technical and empirical observations. First, we need our *k*-mers to be long enough so as to uniquely identify loci in the genome, however not too long so that there is a high chance of errors or truncation during sequencing. Second, we require that these *k*-mers fit in standard integer values using a 2-bit encoding for reasons of efficiency and performance. Thus, we have selected a *k* = 32 for our analysis. We are considering two different types of *k*-mers in Nebula, inner *k*-mers and gapped *k*-mers. We will explain both types below in the context of deletion structural variants.

#### 2.1.1 Inner *k*-mers

Knowing the approximate coordinates of the region affected by a deletion Nebula will query the corresponding reference sequence and create a dictionary of all the *k*-mers of size 32 inside the deletions. As these *k*-mers are extracted from inside the affected sequence, we will refer to them as *inner k-mers*. Nebula will utilize both unique and non-unique (i.e., seen in multiple locations in the reference genome) inner *k*-mers for genotyping. Nebula uses Jellyfish (Marçais and Kingsford (2011)) to generate an index of all the *k*-mers inside the reference genome to help with this step.

As a unique *k*-mer only appears at a single loci in the genome, reads containing it can be assumed to have been generated from sequencing of the same region. Because of this property, the frequency of these *k*-mers in a set of sequencing reads is highly correlated to the copy number of their site-of-origin in the genome of that individual.

Even though a *k*-mer might be non-unique, its different occurrences may still be separable by looking at adjacent *k*-mers in the reference genome. We associate with each *k*-mer a left and right mask which are basically the 32bp sequences surrounding the *k*-mer in the reference assembly. Among all the loci of a *k*-mer, only those with masks similar to those of the loci inside the deleted region will be considered. This idea allows us to effectively reduce the number of loci a non-unique *k*-mer appears at, which can help with the accuracy of the genotyping phase. Note that, a non-unique inner *k*-mer might appear within the boundaries of more than one event. In such cases, all those loci have to be considered when genotyping. The proposed genotyper in the second phase needs to consider the existence of the same *k*-mer in multiple SVs.

#### 2.1.2 Gapped *k*-mers

A deletion (and generally any type of SVs) can result in creation of novel *k*-mers near the breakpoints. As a result, if these *k*-mers are unique, their presence can signal the existence of the deletion. However, it is usually the case that the breakpoints are not exactly known and new *k*-mers may not be reliably constructed. To overcome this problem, we will utilize gapped *k*-mers as defined below.

A gapped *k*-mer, is a sequence of size *k* + *g* from which we discard the middle *g* bases. The resulting gap helps mitigate small inaccuracies in the breakpoint coordinates or *k*-mer sequence. To pick these gapped *k*-mers, a sequence of size 42 spanning over the breakpoint is selected for each event. With the 10 middle bases ignored, left and right halves of 16 bases each are expected to be seen in close proximity in reads from the newly sequenced sample. This process will yield a single gapped *k*-mer per event (Figure 2).

**Figure 2:**
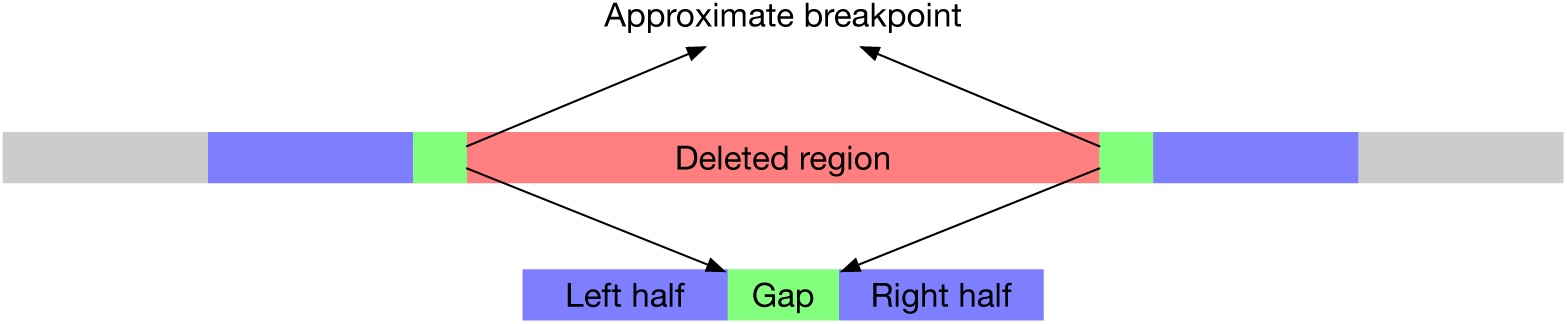
Graphical representation of gapped *k*-mers and their creation.

If the reads from the sample with the alternate allele (i.e., contain the SV) are mapped, the gapped *k*-mers can be extracted from the reads themself by looking at the reads which map near the suspected breakpoints with large soft-clipped sequences. This approach will usually produce few overlapping *k*-mers instead of only one per SV.

### 2.2 Genotyping phase

We formulate the mapping-free genotyping problem using the *k*-mer counts as the optimization problem. The objective is defined as finding the set of genotypes for the SVs of interest which **minimize the total difference between observed** *k***-mer counts and expected values**. There are three main steps to the proposed genotyping phase. First counting the selected *k*-mers in the newly sequenced samples. Second, estimating the optimal real valued relaxed version of the genotypes (0 ≤ *z* ≤ 1). Third, rounding the round the real value relaxed version of genotypes (0 ≤ *z* ≤ 1) into discrete genotypes 0/0, 0/1 or 1/1.

#### 2.2.1 *k*-mer coverage and counting

We define the *k*-mer coverage of a set of sequencing reads as the mean of the number of reads supporting each unique *k*-mer. Assuming the number of reads supporting each *k*-mer follows a normal distribution, the parameters of this distribution can be reasonably estimated by sampling a large number of unique *k*-mers from conserved regions of the reference genome and counting them in the WGS reads of the samples. For more accurate estimates, the GC content of the region surrounding the *k*-mer is also taken into account.

Provided the list of inner and gapped *k*-mers associated with each SV, Nebula proceeds with counting these *k*-mers in the newly sequenced sample. This is done efficiently by treating *k*-mers and sequencing reads as binary values. Expensive string matching operations can now be modeled as lightweight binary arithmetic that translates directly into machine-level instructions. Only those reads containing a *k*-mer surrounded by its masks will be considered towards the final count of that *k*-mer.

#### 2.2.2 Estimating relaxed version of the SV genotypes

Assuming both strands of a genome are sequenced at about the same depth and with similar distribution of reads, count of *k*-mers relative to the sample coverage can act as a proxy for the presence of structural variations. For example, if both copies of a regions are deleted (homozygous deletion), we expect no reads to contain unique *k*-mers coming from that region. On the other hand, a heterozygous deletion will result in inner *k*-mers counts to be around half of the expected count. This observation forms the basis of the proposed genotyping algorithm. Note that, for non-unique *k*-mers Nebula needs to consider the contribution of all those regions in the final count of the non-unique *k*-mers. Same observation holds for gapped *k*-mers, however it is the high count of these *k*-mers that signals the presence of their corresponding events. Nebula can optionally estimate coverage levels for gapped *k*-mers of different gap sizes to possibly improve accuracy at the cost of longer runtime and higher memory usage.

Once both the inner and gapped *k*-mers are selected, Nebula will count them in a single pass of the FASTQ file along with estimating coverage statistics. Given the set of *k*-mers corresponding to an event, the genotype of that event can be deduced by looking at the *k*-mer counts in comparison to the expected *k*-mer counts. We estimate a relaxed version of the genotypes using an objective function of minimizing the difference between observed and expected *k*-mer counts. We utilize linear programming approach for finding the optimal values for this relaxed version of the problem. The objective of the linear program would then be to find the optimal real value relaxation of the genotype for each SV that describes the counts of its *k*-mers.

More formally, given an event *t* with associated *k*-mers *k ∈ K* we define the parameters 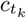, *z*_*t*_, *e*_*k*_ and *C*_*k*_ as follows:

- The variable *z*_*t*_ is the relaxed real value genotype of the event *t* which we would like to calculate.
- Parameter 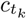 is the expected number of reads that include the *k*-mer *k* produced from the genomic region covered by the event *t* assuming genotype 0/0 for the event *t*.
- *C*_*k*_ is the observed count of *k*-mer *k*
- *e*_*k*_ is an error term for *k*-mer *k* we are minimizing

Assuming the *k*-mer *k* is unique then the basic constraints of the linear programming would be *z_t_* × *c_tk_* + *e_k_* = *C_k_*. Note that, if the *k*-mer *k* appears in more loci in genome than are affected by the events being genotyped we have to update the observed *k*-mer counts *C*_*k*_ to remove expected *k*-mer counts produced from those loci.

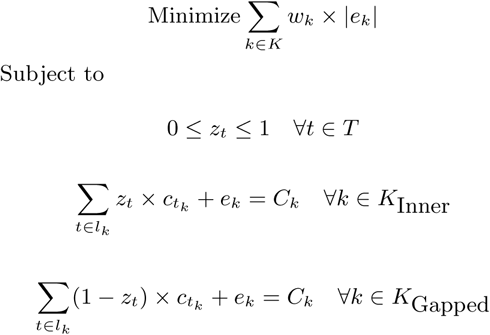

This formulation also supports *k*-mers that appear inside more than one event with *l*_*k*_ being the set of all events which are associated with *k*-mer *k*. Also, as the frequency of gapped and inner *k*-mers follows opposite trends with respect to genotype, the term (1 *-z*_*t*_) has been used for the gapped *k*-mers.

As the number of inner *k*-mers is proportional to the size of the region affected by the event, thus in general there are more inner *k*-mers than gapped *k*-mers associated per SV. However, the gapped *k*-mers tend to yield significantly better genotyping results on their own as compared to inner *k*-mers, suggesting that they are a more reliable signal. As a heuristic, we have add weights to the error terms corresponding to gapped *k*-mers in the objective function of the linear programming. If an event has *i* inner *k*-mers and *g* gapped *k*-mers, its gapped *k*-mers will have a weight of *w*_*k*_ = 2 *× i/g* in the objective function while each inner *k*-mers will have a weight of 1. As events are mostly independent in their set of *k*-mers, the LP can be solved very quickly even for large sets of events and *k*-mers. Nebula relies on CPLEX (IBM (2018)) to generate and solve the linear program.

#### 2.2.3 Rounding

Once the real-valued relaxed version of genotypes (0 ≤ *z*_*t*_ ≤ 1) have been calculated by the linear program for every structural variant *t ∈ T*, a rounding will be applied to decide the final genotypes. Intuitively, a *z*_*t*_ of 1.0 corresponds to a genotype of 0/0, 0.5 corresponds to genotype 1/0 and 0.0 to genotype 1/1. However it may not be clear how to divide the interval [0, 1] between the three genotypes. The naive solution will consider the values *z*_*t*_ falling inside ranges [0, 0.25], [0.25, 0.75] and [0.75, 1] for assigning the SV to genotypes 1/1, 1/0 or 0/0 respectively. However, we here propose a likelihood based approach in picking the rounding ranges.

As mentioned previously, we model a *k*-mer’s count with a normal distribution. If the estimated mean and std of the sample-wide *k*-mer distribution are *σ* and *µ*, then we except the inner unique *k*-mers to follow the below distributions (*b* is a small bias term to avoid a zero standard deviation):

- 𝒩(*µ, σ*^2^) in case of 0/0
- 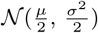 in case of 1/0
- 𝒩(0, *b*) in case of 1/1

We assume for any SV the likelihood of a particular genotype is equal to multiplication of the likelihoods of the counts of its *unique k*-mers coming from that genotype. This can be further simplified to a summation of log-likelihoods as we only care the relative magnitude of likelihoods.

The goal is to divide the interval [0, 1] into three parts by selecting two values *b*_1_ and *b*_2_ for rounding purposes such that events with any *z*_*t*_ falling inside [0, *b*_1_], [*b*_1_, *b*_2_] or [*b*_2_, 1] are considered homozygous, heterozygous or absent respectively. We iterate over all possible pairs of *b*_1_ and *b*_2_ with a step size of 0.01 (under the constraints that *b*_1_ < 0.5 < *b*_2_) and choose the values which yields the highest summation of likelihoods over all *unique k*-mers. After selecting the values *b*_1_ and *b*_2_ we can assign each structural variant *t* ∈ *T* to a specific genotype using the *z*_*t*_ values calculated by LP.

#### 2.2.4 Generalization

The approach described above for deletions can be generalized to support any type of structural variation as long as inner or gapped *k*-mers can be extracted. While certain types of events such as inversions may not produce usable inner *k*-mers, every structural variation can theoretically yield gapped *k*-mers as there is always a breakpoint. The mapping from rounded values to genotypes may need to be reversed depending on whether the event in question reduces or increases the counts of its associated *k*-mers.

## 3 Results

We utilized both simulations and real data to quantify and evaluate the performance of Nebula in genotyping deletions and MEIs reported in the 1000 genomes project Sudmant et al. (2015).

### 3.1 Simulation

We constructed a diploid genome from reference assembly GRCh37 by randomly altering the genome using a total of 4,557 deletion reported on the three samples NA19239, HG00513 and HG00731 in the 1000 genomes project Sudmant et al. (2015). We randomly deleted subset of these 4,557 regions from simulated genome (some deleted from both haplotypes – homozygous (1/1), some deleted from one haplotype – heterozygous (1/0) or not deleted – absent (0/0)). We also considered SNPs at a rate of 1 for every 1000 base pairs in the simulated genome. Then we simulated paired-end sequencing reads from the above genome using `wgsim` (Heng (2011)) with sequencing error rate of 0.001. We considered a high coverage of 40x and a low coverage of 15x for these simulation experiments.

The simulated deletions were used to compare Nebula’s accuracy against those predicted by a combination of Lumpy and SVtyper (Chiang et al. (2015)). First, we compared the false discovery and true discovery rate of Nebula approach versus Lumpy/SVtyper for the 40x simulation. We compared what fraction of deletions were correctly predicted to exist in the simulated genome (i.e., 0/1 or 1/1) and what fraction were correctly predicted to be absent (i.e., 0/0). From 3035 deletions simulated on the genome (i.e., 0/1 or 1/1 genotype) Nebula had a true discovery rate of 0.94 (2855/3035) while Lumpy/Svtyper had a true discovery rate of 0.9 (2747/3035). Furthermore, from the 1522 potential deletions (with genotype 0/0) Nebula had a false discovery rate of 0.03 while Lumpy/SVtyper made no false prediction. Finally, the genotype prediction made by Nebula was comparable to the one produced by SVtyper (Figure 3a) for 40x coverage simulation. Similar results were obtained for the 15x simulation (Figure 3b). Note that, for 198 (*<* 5%) of these deletions Nebula was not able to make any genotype prediction as not enough *k*-mers were found in the preprocessing phase which satisfy the required conditions.

**Figure 3:**
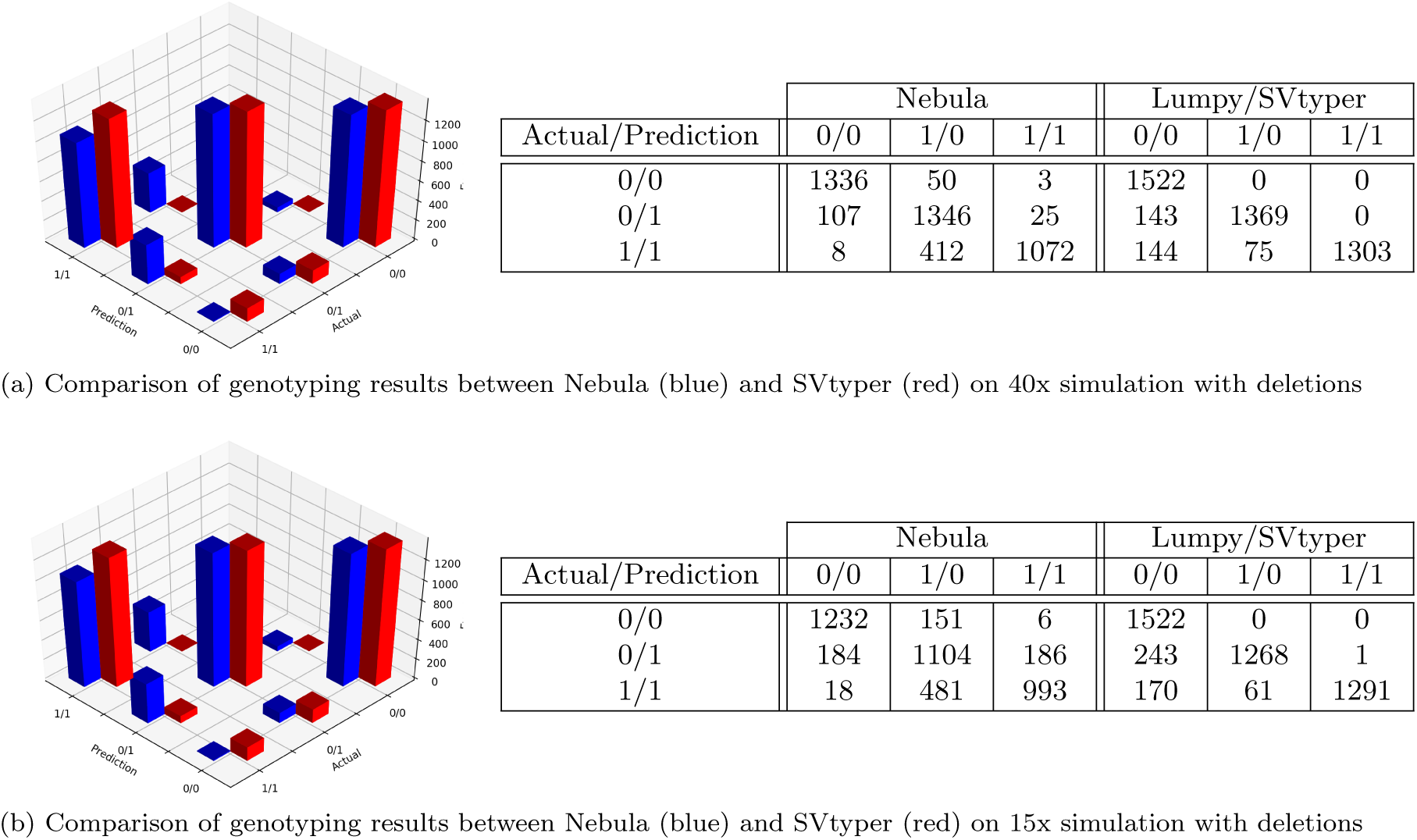
Results of running Nebula to genotype simulated deletions.

### 3.2 Real data

We used the deletions and mobile-element insertions predicted and validated on two samples HG00513 and HG00731 in the 1000 genomes project (Sudmant et al. (2015)) to evaluate performance of Nebula on real WGS data.

We genotyped the deletions reported in sample HG00513 of the 1000 Genomes Project on the WGS data from HG00731 (Figure 4a). Note that, not only the false discovery rate for these deletions was very low (< 5% – the red bars), but also the true discovery rate is quite high (> 95% – the blue bars). When genotyping the heterozygous deletions from HG00731 on HG00513, Nebula reports a homozygous genotype for 176 events. After further investigation we believe that many of these events are truly homozygous deletions and probably incorrectly labeled as heterozygous in previous studies. Similarly, The deletions reported in HG00731 were genotyped using the WGS data from HG00531 (Figure 4b).

**Figure 4:**
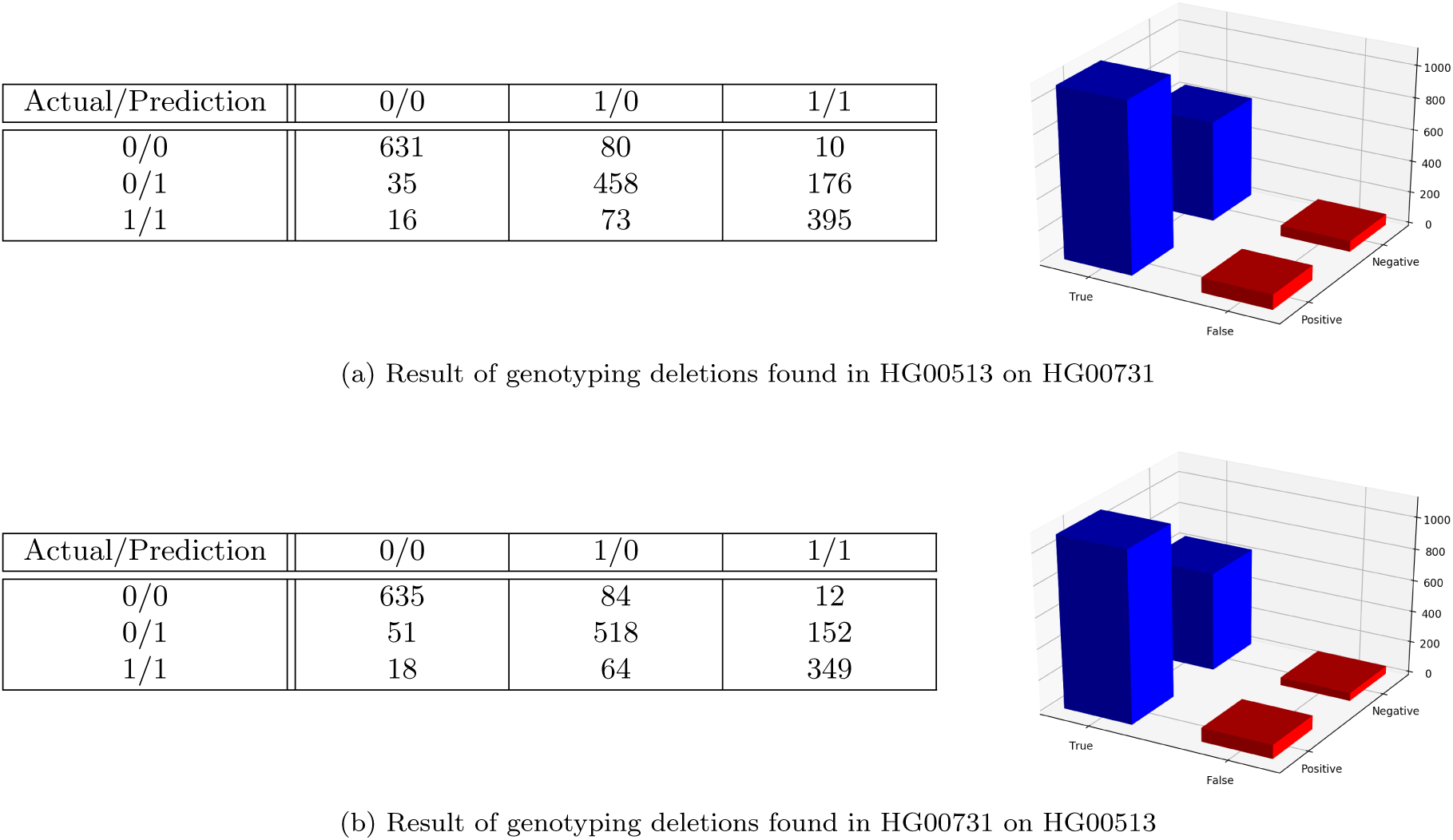
Results of running Nebula to genotype real deletions from HG00513 and HG00731.

A similar procedure was repeated for MEIs reported by 1KG on GRCh38 mapped reads of the same samples. As the original set of events in this case didn’t provide homozygous or heterozygous genotypes, the results have been reported as only present or absent for the sake of consistency. Nebula’s error rate for MEIs remains low at around 3%. These results can be found in Figure 5.

**Figure 5:**
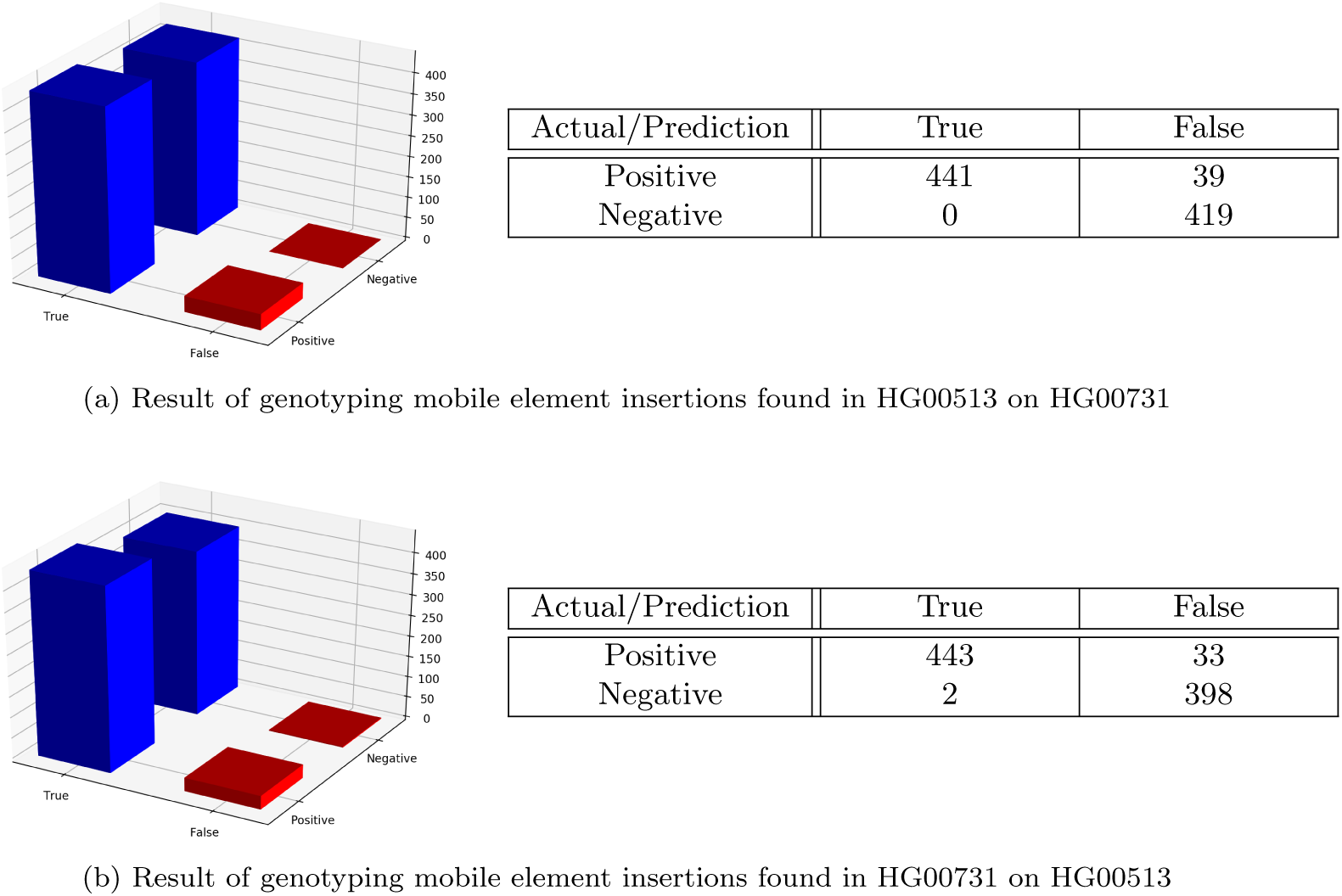
Results of running Nebula to genotype real MEIs from HG00513 and HG00731.

Nebula rarely makes false predictions about the presence or absence of events and a majority of mistakes are classification of homozygous events as heterozygous. Nebula’s false discovery rate for deletions remains at around 8% on real data and as low as 3% on simulation.

Nebula also remains accurate regardless of the size of the event. While shorter events may be harder to genotype due to a lack of inner *k*-mers, the use of gapped *k*-mers assure that such events can be genotyped with reasonable accuracy.

### 3.3 Time and memory performance

The 40x simulation with deletions was also used for runtime and memory usage comparison. All the experiments were run using a single Intel(R) Xeon(R) CPU E5-2670 processor core. Nebula genotyping was compared against the total time required for alignment of sequencing reads using BWA (Li and Durbin (2009); Li (2013)), conversion of reads from SAM format to BAM, sorting and indexing of the BAM file, and the actual runtime of Lumpy/SVtyper (Figure 6).

**Figure 6:**
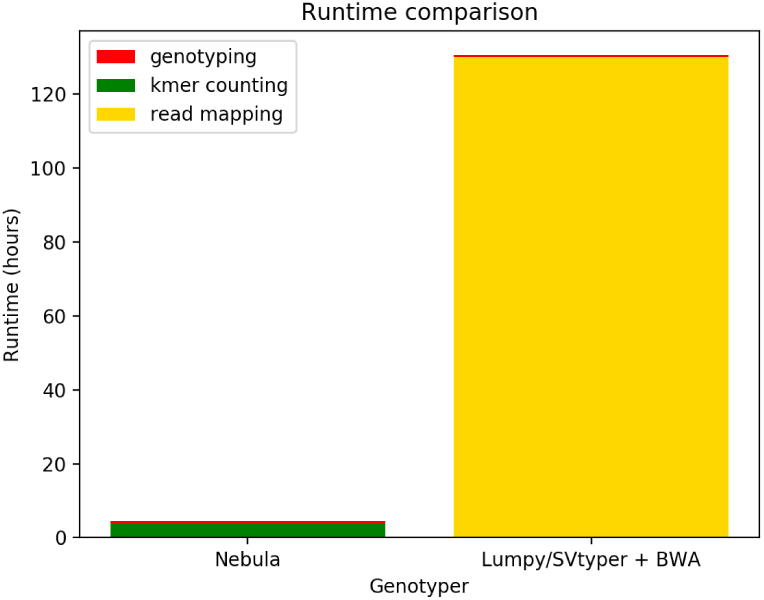
Comparison of runtime of Nebula and BWA+Lumpy/SVtyper on single-core for the 40x simulation.

Due to the entire genotyping process being mapping-free, the runtime of Nebula can be an order of magnitude faster than current approaches. The only time-consuming part of the Nebula’s pipeline is the *k*-mer counting stage. On the simulated sample with nearly 1.8 billion sequencing reads, the counting of nearly half a million *k*-mers (including the gapped and inner *k*-mers of 4500 deletions plus an additional 100000 *k*-mers for estimating coverage statistics) takes slightly less than 4 hours on a single processor core. Once the *k*-mer counts are ready, Nebula’s linear program can be generated and solved in less than 5 minutes (Figure 6).

In comparison, single-core alignment of the same set of reads with BWA takes nearly 110 hours on the same machine. The total run time increases to around 130 hours once all the additional preprocessing required by SVtyper (and Lumpy) are performed. Nebula genotypes the same set of events more than 25 times faster with comparable accuracy.

As we have used a MapReduce (Dean and Ghemawat (2008)) architecture in developing Nebula, the entirety of the pipeline can be accelerated using as many cores as available on a multicore machine. The counting stage of Nebula is particularly IO heavy as it does very minimal processing on each read. As a result, the speed-up of this stage is very dependent on the disk bandwidth available and the runtime may not necessarily decrease linearly with addition of more threads. In our experiments, the *k*-mer counting stage takes nearly 1.5 hours to complete using 4 cores on the same machine, while BWA takes around 28 hours.

Memory usage of Nebula grows almost linearly with the number of events being genotyped. In our simulation, each counter thread requires around 500MB for counting nearly 500000 *k*-mers. As reads are processed on-the-fly, the memory usage remains independent of sample size and is only a function of the number of events being genotyped. BWA on the other hand uses around 5.5GBs of RAM while processing the same sample. It should be noted that due to the MapReduce architecture, *k*-mer count tables are replicated for each counter thread in a copy-on-write fashion, resulting in an almost linear increase of memory usage with addition of threads.

Nebula’s preprocessing stage takes around 4 hours on a single processor core. Selection and filtering of gapped *k*-mers accounts for nearly all the runtime of this stage. The preprocessing step can generate better gapped *k*-mers if mapped reads are available. This will normally be the case as the events in source samples were most likely genotyped using a mapping-based variant caller.

## 4 Conclusion and future work

We’ve demonstrated that *k*-mers can act as a lightweight and simple alternative for expensive mapping when trying to genotype known structural variations. Our approach, Nebula can be used to efficiently genotype these events en masse.

Although we have limited the scope of this paper to genotyping of deletions and MEIs, almost any type of structural variation can theoretically be genotyped by Nebula through the use of gapped *k*-mers extracted from mapped source reads. Inner *k*-mers may not necessarily be available for every type of event, for instance inversions don’t change the frequency of inner *k*-mers as a *k*-mer and its reverse complement are effectively equal in WGS reads.

Furthermore, the proposed approach can easily be extended and parallelized to genotype many samples simultaneously. Thus, we can use a similar approach for genotyping common SVs in large set of WGS cases and controls for finding SVs associated with specific phenotype. Note that, we are currently using a simple *k*-mer counting approach based on standard data structures and hash functions; a more sophisticated approach such as Rizk et al. (2013) could help improve the runtime and reduce the memory usage further.

## 5 Acknowledgements

We would like to acknowledge helpful discussions with Drs. Farhad Hormozdiari, Titus Brown and Daniel Standage regarding this paper.

